# Exploring the phenotypic consequences of tissue specific gene expression variation inferred from GWAS summary statistics

**DOI:** 10.1101/045260

**Authors:** Alvaro N. Barbeira, Scott P. Dickinson, Jason M. Torres, Jiamao Zheng, Eric S. Torstenson, Heather E. Wheeler, Kaanan P. Shah, Rodrigo Bonazzola, Tzintzuni Garcia, Todd Edwards, GTEx Consortium, Dan L. Nicolae, Nancy J. Cox, Hae Kyung Im

**Affiliations:** Section of Genetic Medicine, The University of Chicago, Chicago, IL, USA; Committee on Molecular Metabolism and Nutrition, The University of Chicago, Chicago, IL, USA; Vanderbilt Genetic Institute, Vanderbilt University Medical Center, Nashville, TN, USA; Departments of Biology and Computer Science, Loyola University Chicago, Chicago, IL, USA; Center for Research Informatics, The University of Chicago, IL, USA

## Abstract

Scalable, integrative methods to understand mechanisms that link genetic variants with phenotypes are needed. Here we derive a mathematical expression to compute PrediXcan (a gene mapping approach) results using summary data (S-PrediXcan) and show its accuracy and general robustness to misspecified reference sets. We apply this framework to 44 GTEx tissues and 100+ phenotypes from GWAS and meta-analysis studies, creating a growing public catalog of associations that seeks to capture the effects of gene expression variation on human phenotypes. Replication in an independent cohort is shown. Most of the associations were tissue specific, suggesting context specificity of the trait etiology. Colocalized significant associations in unexpected tissues underscore the need for an agnostic scanning of multiple contexts to improve our ability to detect causal regulatory mechanisms. Monogenic disease genes are enriched among significant associations for related traits, suggesting that smaller alterations of these genes may cause a spectrum of milder phenotypes.

## Introduction

Over the last decade, GWAS have been successful in robustly associating genetic loci to human complex traits. However, the mechanistic understanding of these discoveries is still limited, hampering the translation of the associations into actionable targets. Studies of enrichment of expression quantitative trait loci (eQTLs) among trait-associated variants [1–3] show the importance of gene expression regulation. Functional class quantification showed that 80% of the common variant contribution to phenotype variability in 12 diseases can be attributed to DNAase I hypersensitivity sites, further highlighting the importance of transcript regulation in determining phenotypes [4].

Many transcriptome studies have been conducted where genotypes and expression levels are assayed for a large number of individuals [5–8]. The most comprehensive transcriptome dataset, in terms of examined tissues, is the Genotype-Tissue Expression Project (GTEx): a large-scale effort where DNA and RNA were collected from multiple tissue samples from nearly 1000 individuals and sequenced to high coverage [9, 10]. This remarkable resource provides a comprehensive cross-tissue survey of the functional consequences of genetic variation at the transcript level.

To integrate knowledge generated from these large-scale transcriptome studies and shed light on disease biology, we developed PrediXcan [11], a gene-level association approach that tests the mediating effects of gene expression levels on phenotypes. PrediXcan is implemented on GWAS or sequencing studies (i.e. studies with genome-wide interrogation of DNA variation and phenotypes). It imputes transcriptome levels with models trained in measured transcriptome datasets (e.g. GTEx). These predicted expression levels are then correlated with the phenotype in a gene association test that addresses some of the key limitations of GWAS [11].

Meta-analysis efforts that aggregate results from multiple GWAS have been able to identify an increasing number of associations that were not detected with smaller sample sizes [12–14]. We will refer to these results as GWAMA (Genome-wide association meta-analysis) results. In order to harness the power of these increased sample sizes while keeping the computational burden manageable, methods that use summary level data rather than individual level data are needed.

Methods similar to PrediXcan that estimate the association between intermediate gene expression levels and phenotypes, but use summary statistics have been reported: TWAS (summary version) [15] and SMR (Summary Mendelian Randomization) [16]. Another class of methods that integrate eQTL information with GWAS results are based on colocalization of eQTL and GWAS signals. Colocalized signals provide evidence of possible causal relationship between the target gene of an eQTL and the complex trait. These include RTC [1], Sherlock [17], COLOC [18], and more recently eCAVIAR [19] and ENLOC [20].

Here we derive a mathematical expression that allows us to compute the results of PrediXcan without the need to use individual-level data, greatly expanding its applicability. We compare with existing methods and outline a best practices framework to perform integrative gene mapping studies, which we term MetaXcan.

We apply the MetaXcan framework by first training over 1 million elastic net prediction models of gene expression traits, covering protein coding genes across 44 human tissues from GTEx, and then performing gene-level association tests over 100 phenotypes from 40 large meta-analysis consortia and dbGaP.

## Results

### Computing PrediXcan results using summary statistics

We have derived an analytic expression to compute the outcome of PrediXcan using only summary statistics from genetic association studies. Details of the derivation are shown in the Methods section. In Figure 1-A, we illustrate the mechanics of Summary-PrediXcan (S-PrediXcan) in relation to traditional GWAS and the individual-level PrediXcan method [11].

**Figure 1.**
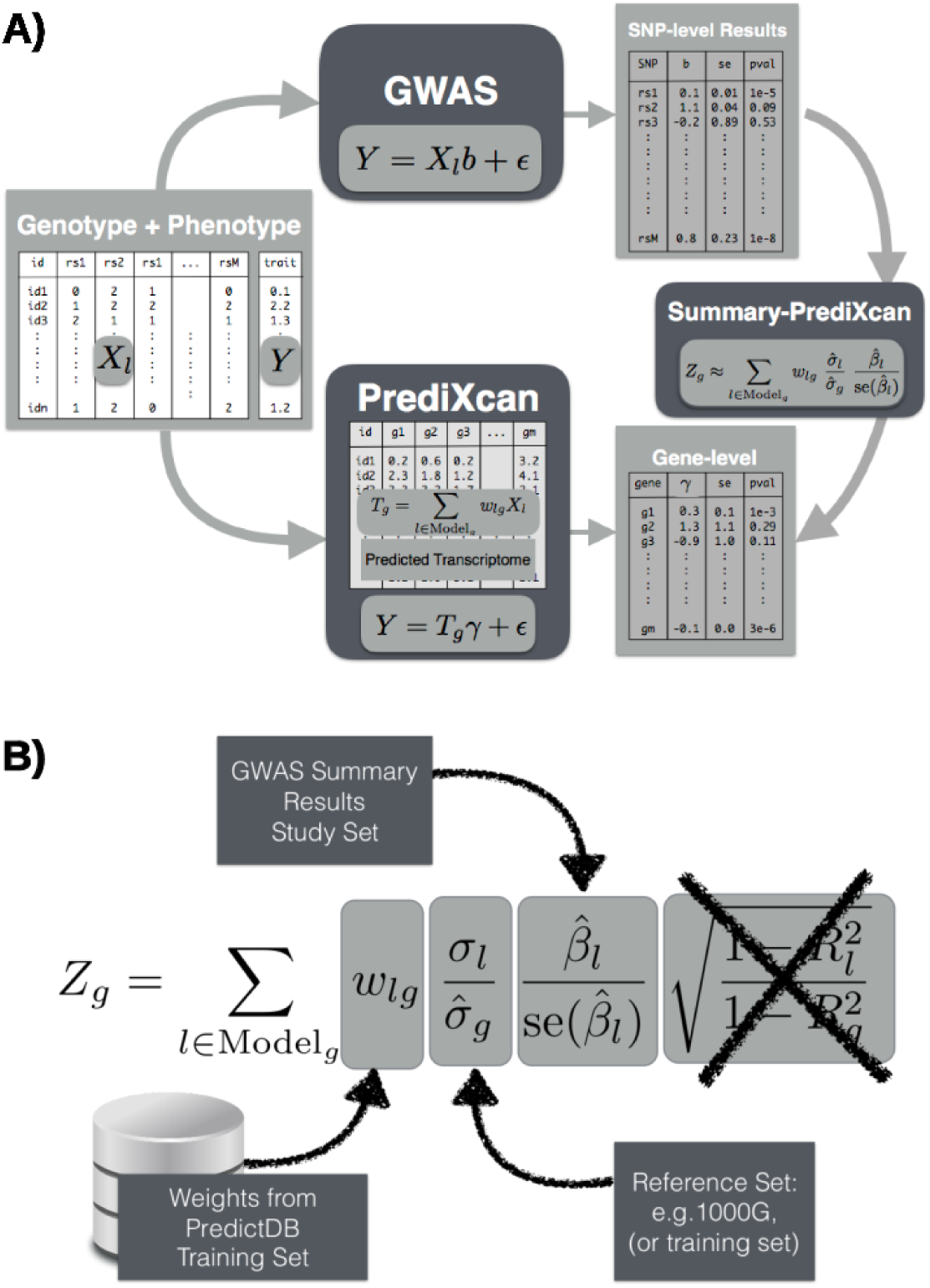
**Panel A: Comparison between GWAS, PrediXcan, and Summary-PrediXcan**. Both GWAS and PrediXcan take genotype and phenotype data as input. GWAS computes the regression coefficients of *Y* on *X*_*l*_ using the model *Y* = *a* + *X*_*l*_*b* + *E*, where *Y* is the phenotype and *X*_*l*_ the individual SNP dosage. The output is a table of SNP-level results. PrediXcan, in contrast, starts first by predicting/imputing the transcriptome. Then it calculates the regression coefficients of the phenotype *Y* on each gene’s predicted expression *T*_*g*_. The output is a table of gene-level results. Summary-PrediXcan directly computes the gene-level association results using the output from GWAS. **Panel B: Components of the S-PrediXcan formula.** This figure shows the components of the formula to calculate PrediXcan gene-level association results using summary statistics. The different sets involved as input data are shown. The regression coefficient between the phenotype and the genotype is obtained from the study set. The training set is the reference transcriptome dataset where the prediction models of gene expression levels are trained. The reference set (1000G, or training set having some advantages) is used to compute the variances and covariances (LD structure) of the markers used in the predicted expression levels. Both the reference set and training set values are pre-computed and provided to the user so that only the study set results need to be provided to the software. The crossed out term was set to 1 as an approximation. We found this approximation to have negligible impact on the results.

We find high concordance between PrediXcan and S-PrediXcan results indicating that in most cases, we can use the summary version without loss of power to detect associations. Figure 2 shows the comparison of PrediXcan and S-PrediXcan Z-scores for a simulated phenotype (under the null hypothesis), a cellular growth phenotype and two disease phenotypes: type 1 diabetes and bipolar disorder from the WTCCC Consortium [21]. For the simulated phenotype, the study sets (in which GWAS is performed) and the reference set (in which LD between SNPs is computed) were African, East Asian, and European from 1000 Genomes. The training set (in which prediction models are trained) was European (DGN Cohort [5]) in all cases. The high correlation between PrediXcan and S-PrediXcan demonstrates the robustness of our method to mismatches between reference and study sets. Despite the generally good concordance between the summary and individual level methods, there were a handful of false positive results with S-PrediXcan much more significant than PrediXcan. This underscores the need to use closely matched LD information whenever possible. Supplementary Figure 11 shows S-PrediXcan’s performance on a phenotype simulated under the alternative hypothesis.

**Figure 2.**
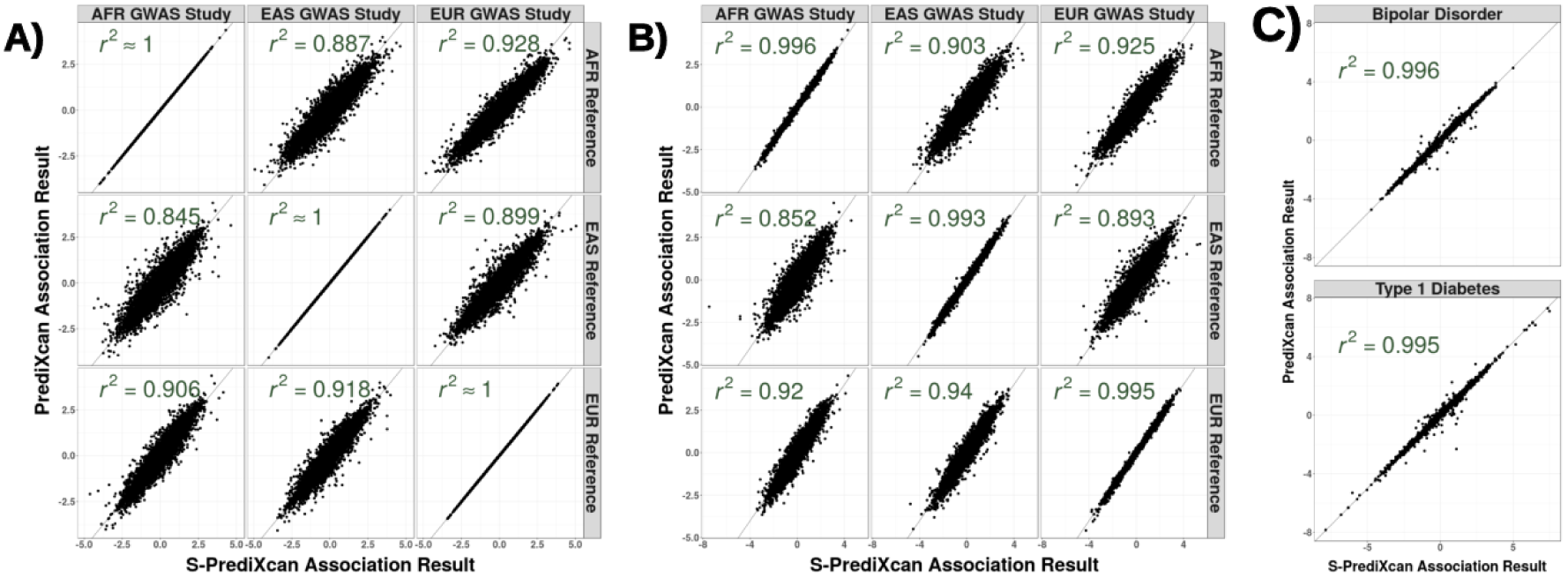
**Comparison of PrediXcan vs. S-PrediXcan** for **A)** a simulated phenotype under null hypothesis of no genetic component; **B)** a cellular phenotype (= intrinsic growth); and **C)** bipolar disorder and type 1 diabetes studies from Wellcome Trust Case Control Consortium (WTCCC). Gene expression prediction models were based on the DGN cohort presented in [11]. For the simulated phenotype, study sets (GWAS set) and reference sets (LD calculation set) consisted of African (661), East Asian (504) and European (503) individuals from the 1000 Genomes Project. When the same study set is used as reference set, we obtained a high correlation: *r*^2^ > 0.99999. For the intrinsic growth phenotype, study sets were a subset of 140 individuals from each of the African, Asian an European groups from 1000 Genomes Project. The reference set was the same as for the simulated phenotype. For the disease phenotypes, the study set consisted of British individuals, and the LD calculation set was the European population subset of the 1000 Genomes Project.

Notice that we are not testing here whether PrediXcan itself is robust to population differences between training and study sets. Robustness of the prediction across populations has been previously reported [22]. We further corroborated this in Supplementary Figure 10.

Next we compare with other summary result-based methods such as S-TWAS, SMR, and COLOC.

### Colocalization estimates complement PrediXcan results

One class of methods seeks to determine whether eQTL and GWAS signals are colocalized or are distinct although linked by LD. This class includes COLOC [18], Sherlock [17], and RTC [1], and more recently eCAVIAR [19], and ENLOC [20]. Thorough comparison between these methods can be found in [18, 19]. HEIDI, the post filtering step in SMR that estimates heterogeneity of GWAS and eQTL signals, can be included in this class. We focus here on COLOC, whose quantification of the probability of five configurations complements well with S-PrediXcan results.

COLOC provides the probability of 5 hypotheses: H0 corresponds to no eQTL and no GWAS association, H1 and H2 correspond to association with eQTL but no GWAS or vice-versa, H3 corresponds to eQTL and GWAS association but independent signals, and finally H4 corresponds to shared eQTL and GWAS association. P0, P1, P2, P3, and P4 are the corresponding probabilities for each configuration. The sum of the five probabilities is 1. The authors [18] recommend to interpret H0, H1, and H2 as limited power; we will aggregate these three hypothesis into one event with probability 1-P3-P4 for convenience.

Figure 3 shows ternary plots [23] with P3, P4, and 1-P3-P4 as vertices. The blue region, top sub-triangle, corresponds to high probability of colocalized eQTL and GWAS signals (P4). The orange region, bottom left, corresponds to high probability of distinct eQTL and GWAS signals (P3). The gray region, center and bottom right, corresponds to low probability of both colocalization and independent signals.

**Figure 3.**
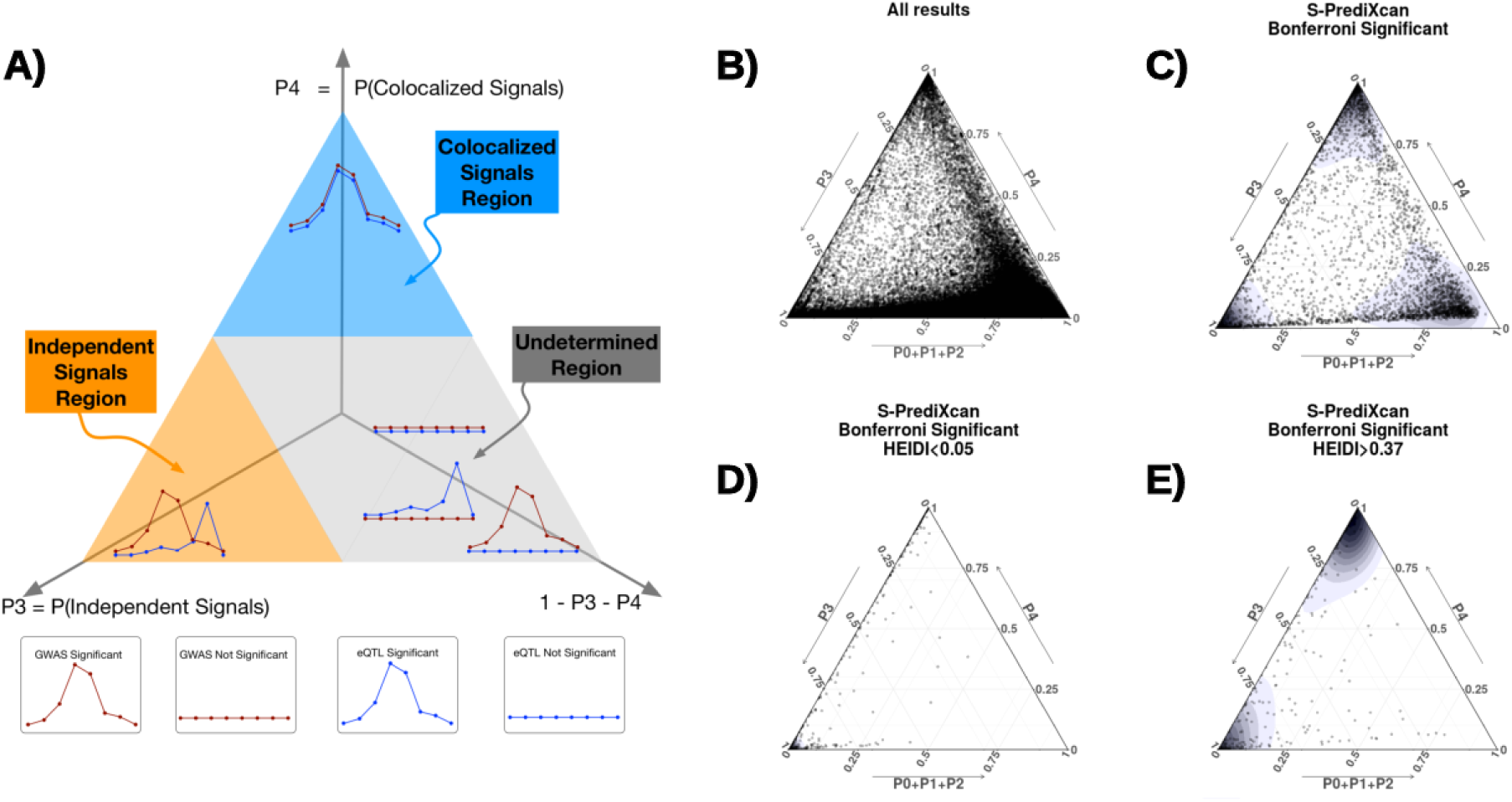
Colocalization status of S-PrediXcan results. **Panel A** shows a ternary plot that represents the probabilities of various configurations from COLOC. This plot conveniently constrains the values such that the sum of the probabilities is 1. All points in a horizontal line have the same probability of ‘colocalized’ GWAS and eQTL signals (P4), points on a line parallel to the right side of the triangle (NW to SE) have the same probability of ‘Independent signals’ (P3), and lines parallel to the left side of the triangle (NE to SW) correspond to constant P1 + P2 + P3. Top sub-triangle in blue corresponds to high probability of colocalization (P4 > 0.5), lower left sub-triangle in orange corresponds to probability of independent signals (P3 > 0.5), and lower right parallelogram corresponds to genes without enough power to determine or reject colocalization. The following panels present ternary plots of COLOC probabilities with a density overlay for S-PrediXcan results of the Height phenotype. **Panel B** shows the colocalization probabilities for all gene-tissue pairs. Most results fall into the ‘undetermined’ region. **Panel C** shows that if we keep only Bonferroni-significant S-PrediXcan results, associations tend to cluster into three distinct regions: ‘independent signals’, ‘colocalized’ and ‘undertermined’. **Panel D** shows that HEIDI significant genes (to be interpreted as high heterogeneity between GWAS and eQTL signals, i.e. distinct signals) tightly cluster in the ‘independent signal’ region, in concordance with COLOC. A few genes fall in the ‘colocalized’ region, in disagreement with COLOC classification. Unlike COLOC results, HEIDI does not partition the genes into distinct clusters and an arbitrary cutoff p-value has to be chosen. **Panel E** shows genes with large HEIDI p-value (no evidence of heterogeneity) which fall in large part in the ‘colocalized’ region. However a substantial number fall in ‘independent signal’ region, disagreeing with COLOC’s classification.

Figure 3-B shows association results for all gene-tissue pairs with the height phenotype. We find that most associations fall in the gray, ‘undetermined’, region. When we restrict the plot to S-PrediXcan Bonferroni-significant genes (Figure 3-C), three distinct peaks emerge in the high P4 region (‘colocalized signals’), high P3 region (‘independent signals’ or ‘non-colocalized signals’), and ‘undetermined’ region. Moreover, when genes with low prediction performance are excluded (Supplementary Figure 6-D) the ‘undetermined’ peak significantly diminishes.

These clusters provide a natural way to classify significant genes and complement S-PrediXcan results. Depending on false positive / false negative trade-off choices, genes in the ‘independent signals’ or both ‘independent signals’ and ‘undetermined’ can be filtered out. The proportion of colocalized associations (P4 > 0.5) ranged from 5 to 100% depending on phenotype with a median of 27.6%. The proportion of ‘non-colocalized’ associations ranged from 0 to 77% with a median of 27.0%. Supplementary Table 2 summarizes the percentages of significant associations that fall into the different colocalization regions.

This post-filtering idea was first implemented in the SMR approach using HEIDI. Comparison of COLOC results with HEIDI is shown in Figure 6-E to-F.

### Comparison of S-PrediXcan to S-TWAS

Gusev et al. have proposed Transcriptome-Wide Association Study based on summary statistics (S-TWAS), which imputes the SNP level Z-scores into gene level Z-scores. This is not the same as computing the results of individual level TWAS. We show (in Methods section) that the difference between the individual level and summary level TWAS is given by the factor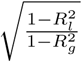, where *R*_*l*_ is the proportion of variance in the phenotype explained by a SNP’s allelic dosage, and *R*_*g*_ is the proportion explained by gene expression (see Methods section). For most practical purposes we have found that this factor is very close to 1 so that if the same prediction models were used, no substantial difference between S-TWAS and S-PrediXcan should be expected.

Figure 4-A shows a diagram of S-PrediXcan and S-TWAS. Both use SNP to phenotype associations results (*Z*_*X*,*Y*_) and prediction weights 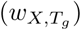 to infer the association between the gene expression level (*T*_*g*_) and phenotype (*Y*).

**Figure 4.**
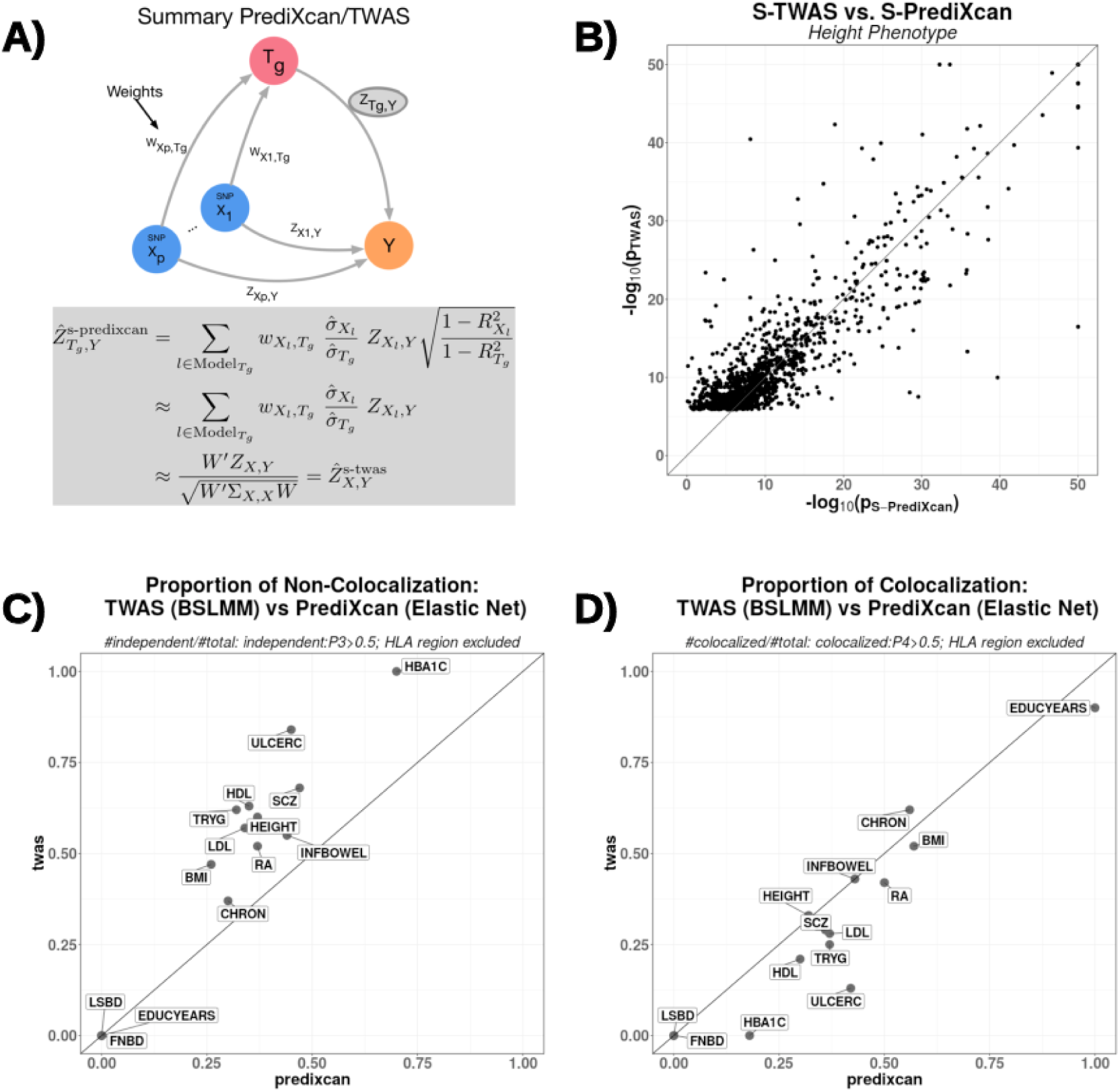
Comparison between S-PrediXcan and S-TWAS. **Panel A** depicts how Summary-TWAS and PrediXcan test the mediating role of gene expression level *T*_*g*_. Multiple SNPs are linked to the expression level of a gene via weights *w*_*X,T*_*g*. **Panel B** shows the significance of Summary-TWAS (BSLMM) vs. Summary-PrediXcan (elastic net), for the height phenotype across 44 GTEx tissues. There is a small bias caused by using S-TWAS results available from [24], which only lists significant hits. S-PrediXcan tends to yield a larger number of significant associations (see Supplementary Figure 13). P-values were thresholded at 10^−50^ for visualization purposes. **Panel C** shows the proportion of non-colocalized associations (distinct eQTL and GWAS signals) from S-TWAS significant vs S-PrediXcan significant results. For all phenotypes, S-TWAS has a higher proportion of LD-contaminated signals compared to S-PrediXcan, as estimated via COLOC. **Panel D** shows the proportion of colocalized associations (shared eQTL and GWAS signals) from S-TWAS significant vs S-PrediXcan significant results. For most phenotypes, TWAS has lower proportion of colocalized signals compared to S-PrediXcan, as estimated via COLOC. Phenotype Abbreviation: Femoral Neck Bone Density (FNBD), Lumbar Spine Bone Density (LSBD), Body Mass Index (BMI), Height (HEIGHT), Low-Density Lipoprotein Cholesterol (LDL), High-Density Lipoprotein Cholesterol (HDL), Tryglicerides (TRYG), Chron’s Disease (CHRON), Inflammatory Bowel’s Disease (INFBOWEL), Ulcerative Colitis (ULCERC), Hemogoblin Levels (HBA1C) HOMA Insulin Response (HOMA-IR) Schizophrenia (SCZ), Rheumatoid Arthritis (RA), College Completion (COLLEGE), Education Years (EDUCYEARS)

Figure 4-B compares S-TWAS significance (as reported in [24]) to S-PrediXcan significance. The difference between the two approaches is mostly driven by the different prediction models: TWAS uses BSLMM [25] whereas PrediXcan uses elastic net [26]. BSLMM allows two components: one sparse (small set of large effect predictors) and one polygenic (all variants contribute some marginal effect to the prediction). For PrediXcan we have chosen to use a sparse model (elastic net) based on the finding that the genetic component of gene expression levels is mostly sparse [27].

Figure 4-C shows that the proportion of non-colocalized (independent) GWAS and eQTL signals is larger among TWAS significant genes than among S-PrediXcan significant ones. This is likely due to the polygenic component of BSLMM models, a wider set of SNPs increasing the chance of capturing LD-contaminated (non-colocalized) association. Figure 4-D shows that, for most traits, the proportion of colocalized signals is larger among S-PrediXcan significant genes than S-TWAS significant genes.

### Comparison of S-PrediXcan to SMR

Zhu et al. have proposed Summary Mendelian Randomization (SMR) [16], a summary data based Mendelian randomization that integrates eQTL results to determine target genes of complex trait-associated GWAS loci. They derive an approximate 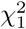-statistic (Eq 5 in [16]) for the mediating effect of the target gene expression on the phenotype.

Unfortunately, the derived statistic is mis-calibrated. A QQ plot comparing the SMR statistic (under the null hypothesis of genome-wide significant eQTL signal and no GWAS association) shows deflation. The sample mean of the statistic is ≈ 0.93 instead of 1, the expected value for the mean of a 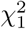 random variable. See Figure 5 (B and C) and Methods section for details. The *χ*^2^ approximation is only valid in two extreme cases: when the eQTL association is much stronger than the GWAS association or vice versa, when the GWAS association is much stronger than the eQTL association. See Methods for details.

**Figure 5.**
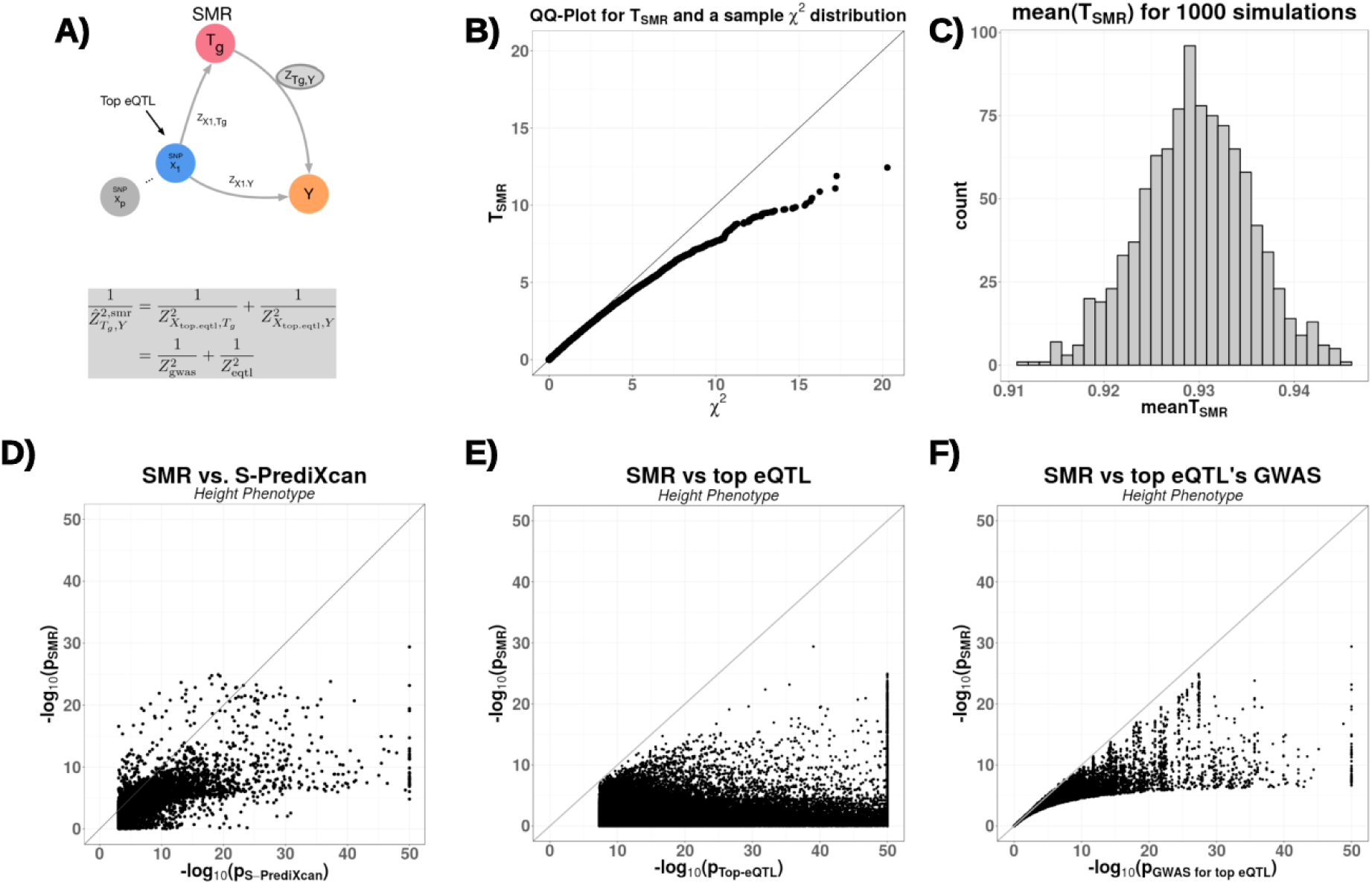
Comparison between Summary-PrediXcan and SMR. **Panel A** depicts how SMR tests the mediating role of gene expression level *T*_*g*_. The top eQTL is linked to the phenotype as an instrumental variable in a Mendelian Randomization approach. **Panel B** shows a QQ plot for simulated values of *T*_SMR_. Under the null hypothesis of significant eQTL signal and no GWAS association, we generated random values for 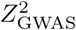 and 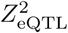 following the simulations from [16]. *T*_SMR_ statistic was calculated from these values, and compared to a 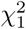 distribution to illustrate this statistics’ deflation. **Panel C** shows the sample mean of *T*_SMR_ from 1000 simulations, centered close to 0.93, instead of the expected value of 1 for a *χ*^2^-distributed variable. **Panel D** shows the significance of SMR vs. the significance of Summary-PrediXcan. As expected, SMR associations tend to be smaller than S-PrediXcan ones. **Panels E** and **F** show that the SMR statistics significance is bounded by GWAS and eQTL p-values. The p-values (- log10) of the SMR statistics are plotted against the GWAS p-value of the top eQTL SNP (**panel E**), and the gene’s top eQTL p-value (**panel F**). Some of the associations, GWAS and eQTL p-values were more significant than shown since they were thresholded at 10^−50^ to improve visualization.

One limitation is that the significance of the SMR statistic is the lower of the top eQTL association (genotype to expression) or the GWAS association (genotype to phenotype) as shown in Figure 5 (E and F). Given the much larger sample sizes of GWAS studies, for most genes, the combined significance will be determined by the eQTL association. The combined statistic forces us to apply multiple testing correction for all genes, even those that are distant to GWAS associated loci, which is unnecessarily conservative. Keep in mind that currently both SMR and PrediXcan only use cis associations. An example may clarify this further. Let us suppose that for a given phenotype there is only one causal SNP and that the GWAS yielded a highly significant p-value, say 10^−50^. Let us also suppose that there is only one gene (gene A) in the vicinity (we are only using cis predictors) associated with the causal SNP with p = 10^−5^. SMR would compute the p-values of all genes and yield a p-value ≈ 10^−5^ for gene A (the less significant p-value). However, after multiple correction this gene would not be significantly associated with the phenotype. Here it is clear that we should not be adjusting for testing of all genes when we know a priori that only one is likely to produce a gene level association. In contrast, the PrediXcan p-value would be ≈ 10^−50^ for gene A and would be distributed uniformly from 0 to 1 for the remaining genes. Most likely only gene A (or perhaps a handful of genes, just by chance) would be significant after Bonferroni correction. If we further correct for prediction uncertainty (here = eQTL association), a p-value of ≈ 10^−5^ would remain significant since we only need to correct for the (at most) handful of genes that were Bonferroni significant for the PrediXcan p-value.

Another potential disadvantage of this method is that only top-eQTLs are used for testing the gene level association. This does not allow to aggregate the effect on the gene across multiple variants.

Figure 5-D compares S-PrediXcan (elastic net) and SMR association results. As expected, SMR p-values tend to be less significant than S-PrediXcan’s in large part due to the additional adjustment for the uncertainty in the eQTL association. Figures 5-E and -F show that the SMR significance is bounded by the eQTL and GWAS association strengths of the top eQTL.

SMR introduces a post filtering step via an approach called HEIDI, which is compared to COLOC in Figure 3 and Supplementary Figure 6.

**Figure 6.**
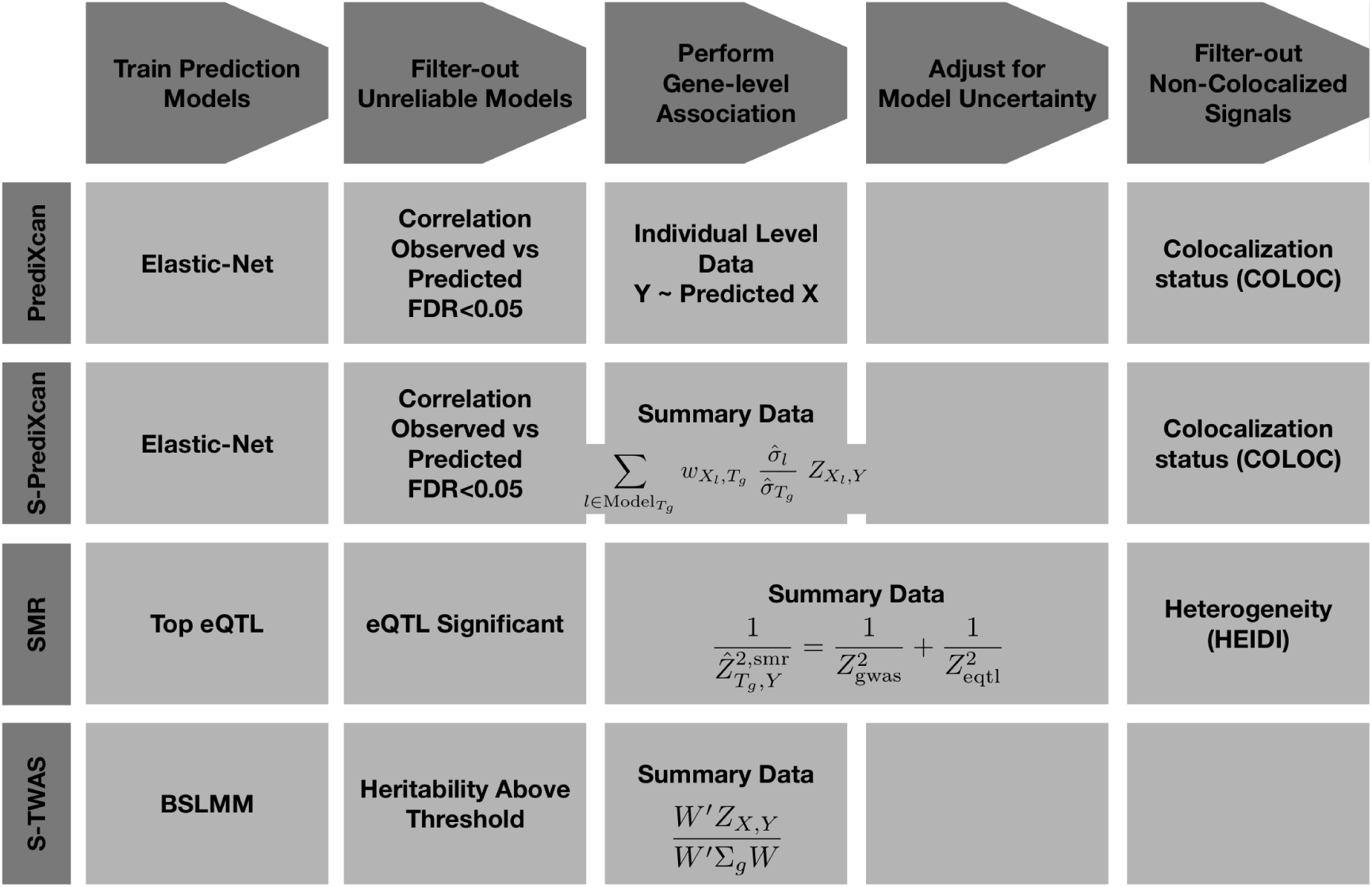
MetaXcan Framework. The figure shows a general framework (MetaXcan) which encompasses methods such as PrediXcan, TWAS, SMR, COLOC among others.

### MetaXcan framework

Building on S-PrediXcan and existing approaches, we define a general framework (MetaXcan) to integrate eQTL information with GWAS results and map disease-associated genes. This evolving framework can incorporate models and methods to increase the power to detect causal genes and filter out false positives. Existing methods fit within this general framework as instances or components (Figure 6).

The framework starts with the training of prediction models for gene expression traits followed by a selection of high-performing models. Next, a mathematical operation is performed to compute the association between each gene and the downstream complex trait. Additional adjustment for the uncertainty in the prediction model can be added. To avoid capturing LD-contaminated associations, which can occur when expression predictor SNPs and phenotype causal SNPs are different but in LD, we use colocalization methods that estimate the probability of shared or independent signals.

PrediXcan implementations use elastic net models motivated by our observation that gene expression variation is mostly driven by sparse components [27]. TWAS implementations have used Bayesian Sparse Linear Mixed Models [25] (BSLMM). SMR fits into this scheme with prediction models consisting solely of the top eQTL for each gene (weights are not necessary here since only one SNP is used at a time).

For the last step, we chose COLOC to estimate the probability of colocalization of GWAS and eQTL signals. COLOC probabilities cluster more distinctly into different classes and thus, unlike other methods, suggests a natural cut off threshold at P = 0.5. Another advantage of COLOC is that for genes with low probability of colocalization, it further distinguishes distinct GWAS and eQTL signals from low power. This is a useful feature that future development of colocalization methods should also offer. SMR, on the other hand, uses its own estimate of ‘heterogeneity’ of signals calculated by HEIDI.

### Suggested association analysis pipeline

1. Perform PrediXcan or S-PrediXcan using all tissues. Use Bonferroni correction for all gene-tissue pairs: keep p < 0.05 / number of gene-tissue pairs tested.
2. Keep associations with significant prediction performance adjusting for number of PrediXcan significant gene-tissue pairs: keep prediction performance p-values < 0.05 / (number of significant associations from previous step).
3. Filter out LD-contaminated associations, i.e. gene-tissue pairs in the ‘independent signal’ (=‘non-colocalized’) region of the ternary plot (See Figure 3-A): keep COLOC P3 < 0.5 (Blue and gray regions in Figure 3-A).
4. If further reduction of number of genes to be taken to replication or validation is desired, keep only hits with explicit evidence of colocalization: P4 > 0.5 (Blue region in Figure 3-A).

Any choice of thresholds has some level of arbitrariness. Depending on the false positive and negative trade off, these number may be changed.

### Gene expression variation in humans is associated to diverse phenotypes

We downloaded summary statistics from meta analyses of over 100 phenotypes from 40 consortia. The full list of consortia and phenotypes is shown in Supplementary Table 4. We tested association between these phenotypes and the predicted expression levels using elastic net models in 44 human tissues from GTEx as described in the Methods section, and a whole blood model from the DGN cohort presented in [11].

We used a Bonferroni threshold accounting for all the gene-tissue pairs that were tested (0.05 / total number of gene-tissue pairs ≈ 2.5e-7). This approach is conservative because the correlation between tissues would make the total number of independent tests smaller than the total number of gene-tissue pairs. Height had the largest number of significantly associated unique genes at 1,686 (based on a GWAMA of 250K individuals). Other polygenic diseases with a large number of associations include schizophrenia with 305 unique significant genes (*n* = 150K individuals), low-density lipoprotein cholesterol (LDL-C) levels with 296 unique significant genes (*n* = 188K), other lipid levels, glycemic traits, and immune / inflammatory disorders such as rheumatoid arthritis and inflammatory bowel disease. For other psychiatric phenotypes, a much smaller number of significant associations was found, with 8 significant genes for bipolar disorder (*n* = 16,731) and one for major depressive disorder (*n* = 18,759), probably due to smaller sample sizes, but also smaller effect sizes.

When step 2 from the suggested pipeline is applied, keeping only reliably predicted genes, we are left with 739 genes for height, 150 for schizophrenia, 117 for LDL-C levels.

After step 3, which keeps genes that are without strong evidence of LD-contamination, these numbers dropped to 264 for height, 58 for schizophrenia, and 60 for LDL-C levels. After step 4, which keeps only genes with strong evidence of colocalization, we find 215 genes for height, 49 for schizophrenia, and 35 for LDL-C. The counts for the full set of phenotypes can be found in Supplementary Table 4.

Mostly, genome-wide significant genes tend to cluster around known SNP-level genome-wide significant loci or sub-genome-wide significant loci. Regions with sub-genome-wide significant SNPs can yield genome-wide significant results in S-PrediXcan, because of the reduction in multiple testing and the increase in power arising from the combined effects of multiple variants. Supplementary Table 3 lists a few examples where this occurs.

The proportion of colocalized associations (P4 > 0.5) ranged from 5 to 100% depending on phenotype with a median of 27.6%. The proportion of ‘non colocalized’ associations ranged from 0 to 77% with a median of 27.0%.

See full set of results in our online catalog (gene2pheno.org). Significant gene-tissue pairs are included in Supplementary Table 5. To facilitate comparison, the catalog contains all SMR results we generated and the S-TWAS results reported by [24] for 30 GWAS traits and GTEx BSLMM models. Note that SMR application to 28 phenotypes was reported by [28] using whole blood eQTL results from [29].

### Moderate changes in ClinVar gene expression is associated with milder phenotypes

We reasoned that if complete knock out of monogenic disease genes cause severe forms of the disease, more moderate alterations of gene expression levels (as effected by regulatory variation in the population) could cause more moderate forms of the disease. Thus moderate alterations in expression levels of monogenic disease genes (such as those driven by eQTLs) may have an effect on related complex traits, and this effect could be captured by S-PrediXcan association statistics. To test this hypothesis, we obtained genes listed in the ClinVar database [30] for obesity, rheumatoid arthritis, diabetes, Alzheimer’s, Crohn’s disease, ulcerative colitis, age-related macular degeneration, schizophrenia, and autism. Figure 8 displays the QQ plot for all associations and those in ClinVar database. As postulated, we found enrichment of significant S-PrediXcan associations for ClinVar genes for all tested phenotypes except for autism and schizophrenia. The lack of significance for autism is probably due to insufficient power: the distribution of p-values is close to the null distribution. In contrast, for schizophrenia, many genes were found to be significant in the S-PrediXcan analysis. There are several reasons that may explain this lack of enrichment: genes identified with GWAS and subsequently with S-PrediXcan have rather small effect sizes, so that it would not be surprising that they were missed until very large sample sizes were aggregated; ClinVar genes may originate from rare mutations that are not well covered by our prediction models, which are based on common variation (due to limited sample sizes of eQTL studies and the minor allele frequency-MAF-filter used in GWAS studies); or the mechanism of action of the schizophrenia linked ClinVar genes may be different than the alteration of expression levels. Also, the pathogenicity of some of the ClinVar entries has been questioned [31]. The list of diseases in ClinVar used to generate the enrichment figures can be found in Supplementary Table 1, along with the corresponding association results.

**Table 1.** Replication of results in GERA. Significant genes / tissue pairs were replicated using a closely matched phenotype in an independent dataset from the GERA cohort [36]. The criteria consisted in significance threshold for replication at p < 0.05, concordant directions of effect, and meta analysis p-value less than the Bonferroni threshold in the discovery set. *π*_1_ is an estimate of proportion of true positives in the replication set. *π*_1_(all) uses all gene-tissue pairs whereas *π*_1_(sig) is computed using only gene-tissue pairs that were significant in the discovery set. The column ‘# replicated genes coloc or undeterm’ is the number of replicated genes excluding the ones for which there was strong evidence of independent GWAS and eQTL signals.

| Discovery phenotype | Replication phenotype | # signif genes in disc set | # replicated genes | $\pi_1$(all) in repl | $\pi_1$(sig) in repl | % replicated genes | # replicated coloc or undeterm |
| --- | --- | --- | --- | --- | --- | --- | --- |
| Coronary artery disease | Any cardiac event | 56 | 6 | 0.4% | 49.1% | 10.7% | 6 |
| LDL cholesterol | Dyslipidemia | 282 | 219 | 5.8% | 90.8% | 78.5% | 184 |
| Triglycerides | Dyslipidemia | 233 | 100 | 5.8% | 73.1% | 43.5% | 69 |
| Schizophrenia | Any psychiatric event | 285 | 60 | 1.2% | 47.6% | 21.1% | 51 |

**Figure 7.**
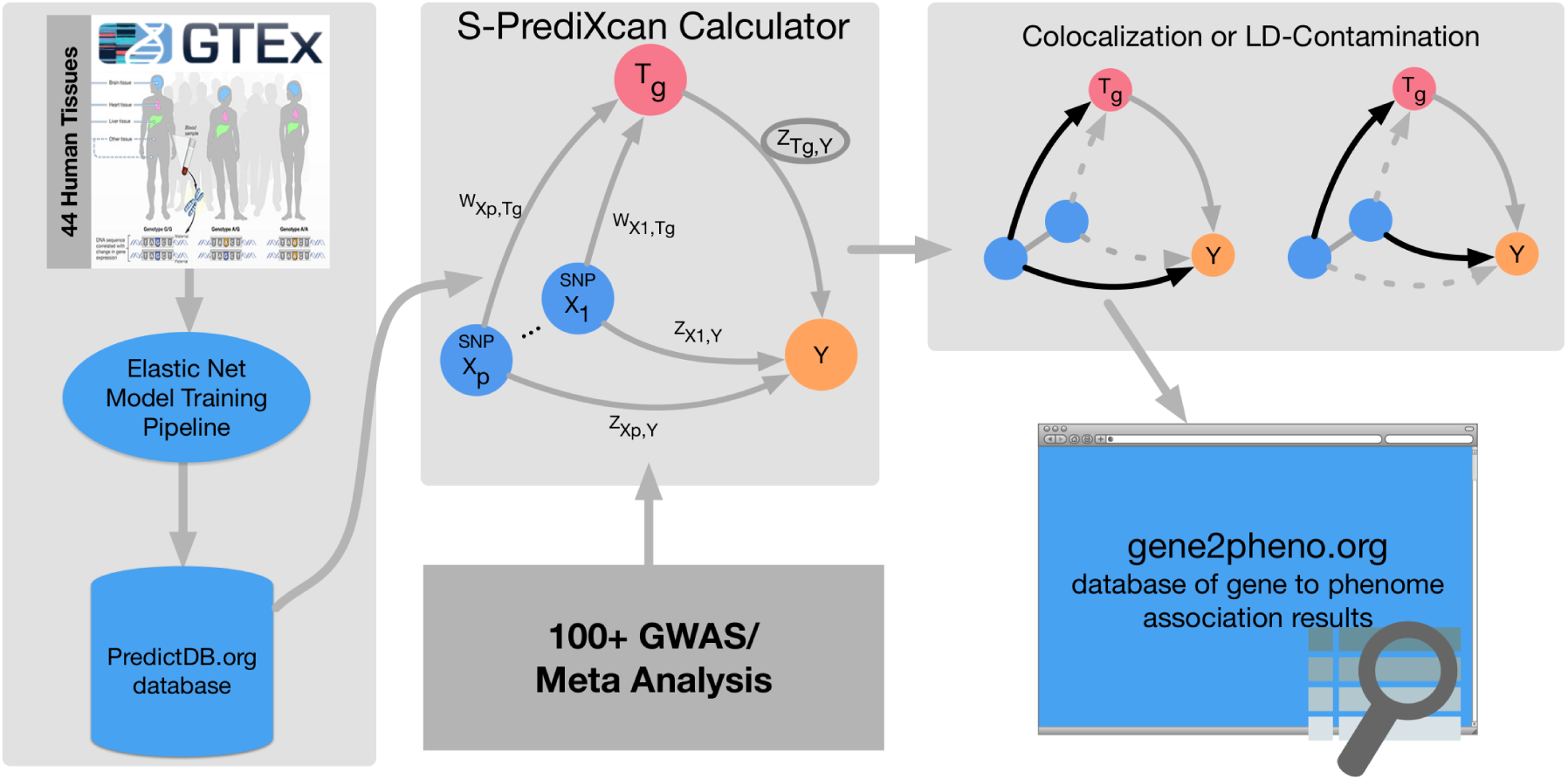
MetaXcan Framework application. This figure summarizes the application of the MetaXcan framework with S-PrediXcan using 44 GTEx tissue transcriptomes and over 100 GWAS and meta analysis results. We trained prediction models using elastic-net [26] and deposited the weights and SNP covariances in the publicly available resource (http://predictdb.org/). The weights, covariances and over 100 GWAS summary results were processed with S-PrediXcan. Colocalization status was computed and the full set of results was deposited in gene2pheno.org.

**Figure 8.**
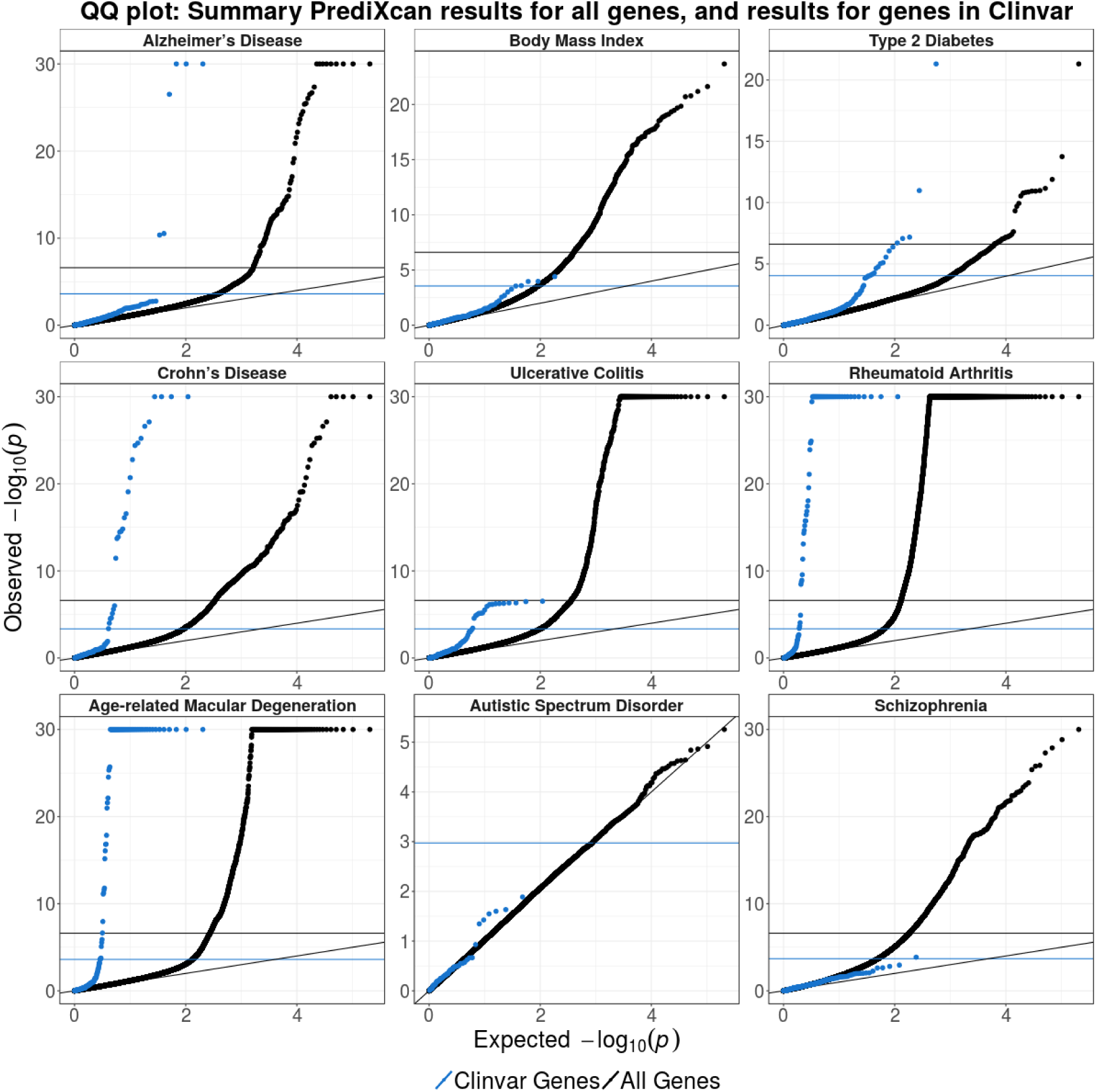
ClinVar genes show significant S-PrediXcan associations. Genes implicated in ClinVar tended to be more significant in S-PrediXcan for most diseases tested, except for schizophrenia and autism. This suggests that more moderate alteration of monogenic disease genes may contribute in a continuum of more moderate but related phenotypes. Alternatively, a more complex interplay between common and rare variation could be taking place such as higher tolerance to loss of function mutations in lower expressing haplotypes which could induce association with predicted expression. Blue circles correspond to the QQ plot of genes in ClinVar that were annotated with the phenotype and black circles correspond to all genes.

### Agnostic scanning of a broad set of tissues enabled by GTEx improves discovery

Most genes were found to be significantly associated in a handful of tissues as illustrated in Figure 9-B. For example, for LDL-C levels, liver was the most enriched tissue in significant associations as expected given known biology of this trait. (See Supplementary Figure 5). This prominent role of liver was apparent despite the smaller sample size available for building liver models (n = 97), which was less than a third of the numbers available for muscle (n = 361) or lung (n = 278).

**Figure 9.**
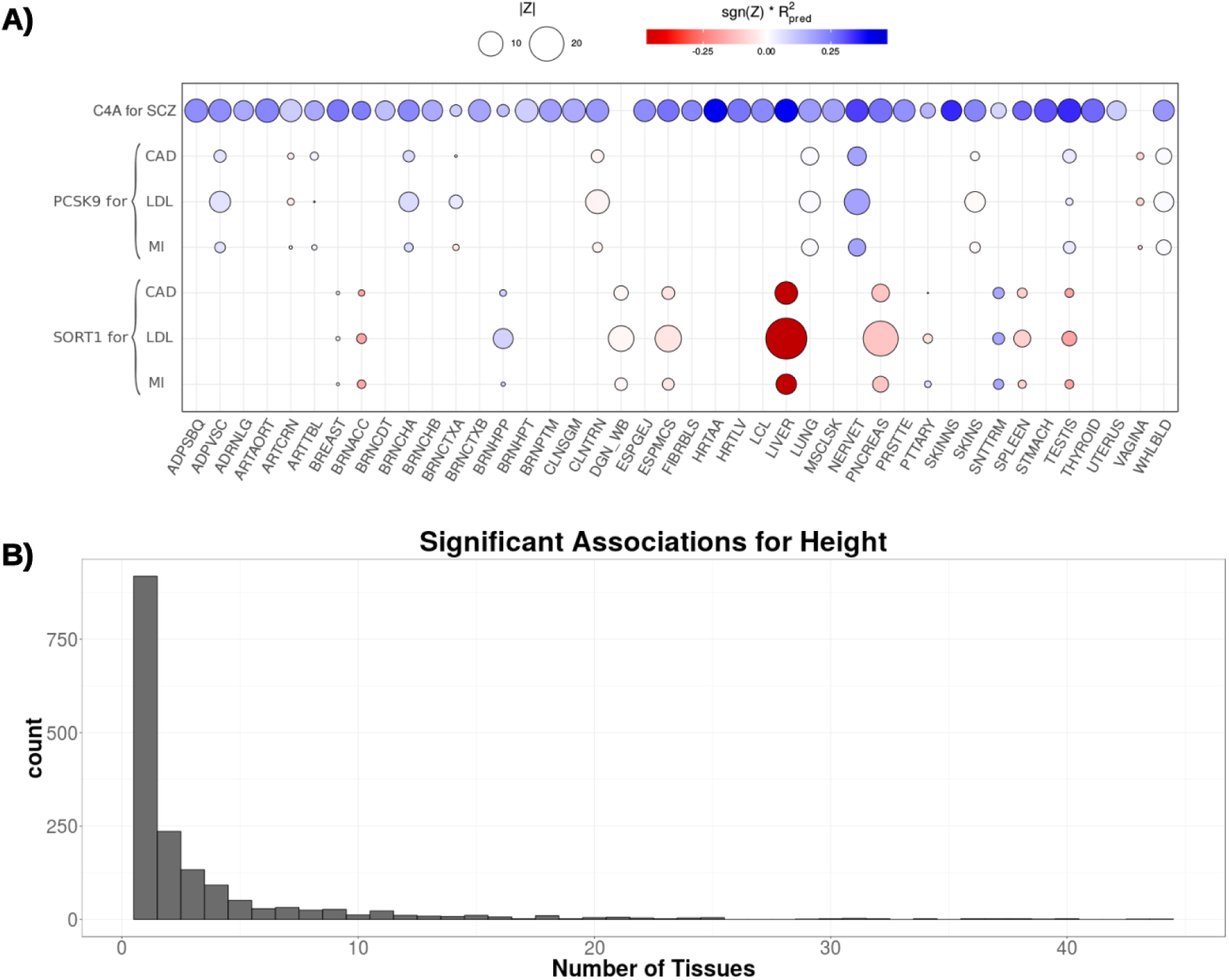
A) S-PrediXcan association for *PCSK9*, *SORT1*, and *C4A* by tissue. This figure shows the association strength between three well studied genes and corresponding phenotypes. *C4A* associations with schizophrenia (SCZ) are significant across most tissues. *SORT1* associations with LDL-C, coronary artery disease (CAD), and myocardial infarction (MI) are most significant in liver. *PCSK9* associations with LDL-C, coronary artery disease (CAD), and myocardial infarction (MI) are most significant in tibial nerve. The size of the points represent the significance of the association between predicted expression and the traits indicated on the top labels. Red indicates negative correlation whereas blue indicates positive correlation. 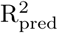 is a performance measure computed as the correlation squared between observed and predicted expression, cross validated in the training set. Darker points indicate larger genetic component and consequently more active regulation in the tissue. **B) Tissue specificity of most trait associations.** This figure shows a histogram of the number of tissues for which a gene is significantly associated with height (other phenotypes show similar pattern). Tissue abbreviation: Adipose-Subcutaneous (ADPSBQ), Adipose-Visceral (Omentum) (ADPVSC), Adrenal Gland (ADRNLG), Artery – Aorta (ARTAORT), Artery-Coronary (ARTCRN), Artery-Tibial (ARTTBL), Bladder (BLDDER), Brain-Amygdala (BRNAMY), Brain-Anterior cingulate cortex (BA24) (BRNACC), Brain-Caudate (basal ganglia) (BRNCDT), Brain-Cerebellar Hemisphere (BRNCHB), Brain-Cerebellum (BRNCHA), Brain-Cortex (BRNCTXA), Brain-Frontal Cortex (BA9) (BRNCTXB), Brain-Hippocampus (BRNHPP), Brain-Hypothalamus (BRNHPT), Brain-Nucleus accumbens (basal ganglia) (BRNNCC), Brain-Putamen (basal ganglia) (BRNPTM), Brain-Spinal cord (cervical c-1) (BRNSPC), Brain-Substantia nigra (BRNSNG), Breast-Mammary Tissue (BREAST), Cells-EBV-transformed lymphocytes (LCL), Cells-Transformed fibroblasts (FIBRBLS), Cervix-Ectocervix (CVXECT), Cervix-Endocervix (CVSEND), Colon-Sigmoid (CLNSGM), Colon-Transverse (CLNTRN), Esophagus Gastroesophageal Junction (ESPGEJ), Esophagus-Mucosa (ESPMCS), Esophagus-Muscularis (ESPMSL), Fallopian Tube (FLLPNT), Heart-Atrial Appendage (HRTAA), Heart-Left Ventricle (HRTLV), Kidney-Cortex (KDNCTX), Liver (LIVER), Lung (LUNG), Minor Salivary Gland (SLVRYG), Muscle-Skeletal (MSCLSK), Nerve-Tibial (NERVET), Ovary (OVARY), Pancreas (PNCREAS), Pituitary (PTTARY), Prostate (PRSTTE), Skin-Not Sun Exposed (Suprapubic) (SKINNS), Skin-Sun Exposed (Lower leg) (SKINS), Small Intestine-Terminal Ileum (SNTTRM), Spleen (SPLEEN), Stomach (STMACH), Testis (TESTIS), Thyroid (THYROID), Uterus (UTERUS), Vagina (VAGINA), Whole Blood (WHLBLD).

However, in general, tissues expected to stand out as more enriched for diseases given currently known biology did not consistently do so when we looked at the average across all (significant) genes, using various measures of enrichment. For example, the enrichment in liver was less apparent for high-density lipoprotein cholesterol (HDL-C) or triglyceride levels. We find for many significant associations that the evidence is present across multiple tissues. This may be caused by a combination of context specificity and sharing of regulatory mechanism across tissues.

Next, we illustrate the challenges of identifying disease relevant tissues based on eQTL information using three genes with well established biology: *C4A* for schizophrenia [32] and *SORT1* [33] and *PCSK9* both for LDL-C and cardiovascular disease. S-PrediXcan results for these genes and traits, and regulatory activity by tissue (as measured by the proportion of expression explained by the genetic component), are shown in Figure 9-A. Representative results are shown in Supplementary Tables 6, 7, and 8. Supplementary Table 9 contains the full set MetaXcan results (i.e. association, colocalization, HEIDI) for these genes.

*SORT1* is a gene with strong evidence for a causal role in LDL-C levels, and as a consequence, is likely to affect risk for cardiovascular disease [33]. This gene is most actively regulated in liver (close to 50% of the expression level of this gene is determined by the genetic component) with the most significant S-PrediXcan association in liver (p-value ≈ 0, *Z* = −28.8), consistent with our prior knowledge of lipid metabolism. In this example, tissue specific results suggest a causal role of *SORT1* in liver.

However, in the following example, association results across multiple tissues do not allow us to discriminate the tissue of action. *C4A* is a gene with strong evidence of causal effect on schizophrenia risk via excessive synaptic pruning in the brain during development [32]. Our results show that *C4A* is associated with schizophrenia risk in all tissues (p < 2.5 *×* 10^−7^ in 36 tissue models and p < 0.05 for the remaining 4 tissue models).

*PCSK9* is a target of several LDL-C lowering drugs currently under trial to reduce cardiovascular events [34]. The STARNET study [35] profiled gene expression levels in cardiometabolic disease patients and showed tag SNP rs12740374 to be a strong eQTL for *PCSK9* in visceral fat but not in liver. Consistent with this, our S-PrediXcan results also show a highly significant association between *PCSK9* and LDL-C (p ≈ 10^−13^) in visceral fat and not in liver (our training algorithm did not yield a prediction model for *PCSK9*, i.e. there was no evidence of regulatory activity). In our results, however, the statistical evidence is much stronger in tibial nerve (p ≈ 10^−27^). Accordingly, in our training set (GTEx), there is much stronger evidence of regulation of this gene in tibial nerve compared to visceral fat.

Most associations highlighted here have high colocalization probabilities. See Supplementary tables 6, 7, and 8. However, visceral fat association shows evidence of non colocalization (probability of independent signals P3 = 0.69 in LDL-C). It is possible that the relevant regulatory activity in visceral adipose tissue was not detected in the GTEx samples for various reasons but it was detected in tibial nerve. Thus by looking into all tissues’ results we increase the window of opportunities where we can detect the association.

*PCSK9* yields colocalized signals for LDL-C levels in Tibial Nerve, Lung and Whole blood. *SORT1* shows colocalization with LDL-C in liver (P4 ≈ 1) and pancreas (P4 = 0.90). *C4A* is colocalized with schizophrenia risk for the majority of the tissues (29 / 40) with a median colocalization probability of 0.82.

These examples demonstrate the power of studying regulation in a broad set of tissues and contexts and emphasize the challenges of determining causal tissues of complex traits based on in-silico analysis alone. Based on these results, we recommend to scan all tissues’ models to increase the chances to detect the relevant regulatory mechanism that mediates the phenotypic association. False positives can be controlled by Bonferroni correcting for the additional tests.

### Replication in an independent cohort, GERA

We used data from the Resource for Genetic Epidemiology Research on Adult Health and Aging study (GERA, phs000674.v1.p1) [36, 37]. This is a study led by the Kaiser Permanente Research Program on Genes, Environment, and Health (RPGEH) and the UCSF Institute for Human Genetics with over 100,000 participants. We downloaded the data from dbGaP and performed GWAS followed by S-PrediXcan analysis of 22 conditions available in the European subset of the cohort.

For replication, we chose Coronary Artery Disease (CAD), LDL cholesterol levels, Triglyceride levels, and schizophrenia, which had closely related phenotypes in the GERA study and had a sufficiently large number of Bonferroni significant associations in the discovery set. Analysis and replication of the type 2 diabetes phenotype can be found in [38]. Coronary artery disease hits were compared with ‘Any cardiac event’, LDL cholesterol and triglyceride level signals were compared with ‘Dyslipidemia’, and schizophrenia was compared to ‘Any psychiatric event’ in GERA.

High concordance between discovery and replication is shown in Figure 10 where dyslipidemia association Z-scores are compared to LDL cholesterol Z-scores. The majority of gene-tissue pairs (92%, among the ones with Z-score magnitude greater than 2 in both sets) have concordant direction of effects in the discovery and replication sets. The high level of concordance is supportive of an omnigenic trait architecture [39]

**Figure 10.**
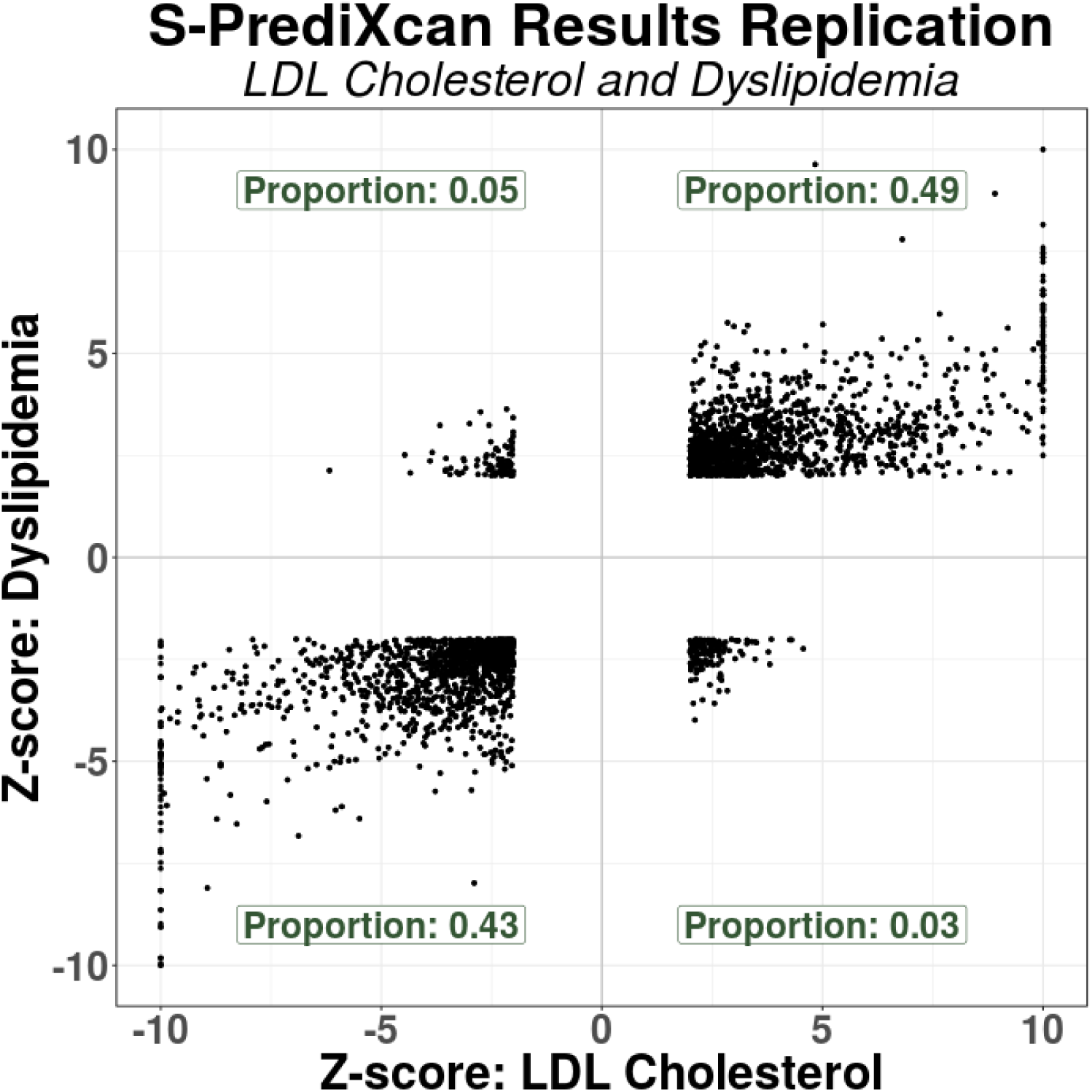
Discovery and replication Z-scores for lipid trait. This figure shows the Z-scores of the association between dyslipidemia (GERA) and predicted gene expression levels on the vertical axis and the Z-scores for LDL cholesterol on the horizontal axis. To facilitate visualization, very large Z-scores where thresholded to 10. Proportions in each quadrant were computed excluding Z-scores with magnitude smaller than 2 to filter out noise.

Following standard practice in meta-analysis, we consider a gene to be replicated when the following three conditions are met: the p-value in the replication set is < 0.05, the direction of discovery and replication effects are the same, and the meta analyzed p-value is Bonferroni significant with the discovery threshold.

Among the 56 genes significantly associated with CAD in the discovery set, 6 (11%) were significantly associated with ‘Any cardiac event’ in GERA. Using ‘Dyslipidemia’ as the closest matching phenotype, 78.5% and 43.5% of LDL and triglyceride genes replicated, respectively. Among the 285 genes associated with schizophrenia in the discovery set, 51 (21%) replicated. The low replication rate for CAD and Schizophrenia is likely due to the broad phenotype definitions in the replication.

We found no consistent replication pattern difference between colocalized and non-colocalized genes. This is not unexpected if the LD pattern is similar between discovery and replication sets.

The full list of significant genes can be queried in gene2pheno.org.

## Discussion

Here we derive a mathematical expression to compute PrediXcan results without using individual level data, which greatly expands its applicability and is robust to study and reference set mismatches. This has not been done before. TWAS, which for the individual level approach only differs from PrediXcan on the prediction model used in the implementation, has been extended to use summary level data. When Gaussian imputation is used, the relationship between individual level and summary versions of TWAS is clear. This is not the case when extended to general weights (such as BSLMM). Our mathematical derivation shows the analytic difference between them explicitly.

The larger proportion of non-colocalized signals from TWAS suggests that by using BSLMM (with polygenic component), it could be more susceptible to LD-contamination than PrediXcan, which uses elastic net (a sparser model). Improved colocalization methods without single causal variant assumption may be needed to strengthen this argument. But the predominantly sparse genetic architecture of gene expression traits [27] supports the benefit of elastic net over BSLMM predictors. We show that SMR statistics needs to be calibrated and argue that by combining the eQTL and GWAS uncertainties into one statistic, it forces the user to apply multiple correction that may be unnecessarily conservative.

We also add a post filtering step, to mitigate issues with LD-contamination. Based on consistency with PrediXcan and interpretability of results, we have chosen to use COLOC for filtering. However, colocalization estimation is an active area of research and improved versions or methods will be adopted in the future.

Despite the generally good concordance between the summary and individual level methods, there were a handful of false positive results with S-PrediXcan much more significant than PrediXcan. This underscores the need to use closely matched LD information whenever possible.

We applied our framework to over 100 phenotypes using transcriptome prediction models trained in 44 tissue from the GTEx Consortium and generated a catalog of downstream phenotypic association results of gene expression variation, a growing resource for the community.

The enrichment of monogenic disease genes among related phenotype associations suggests that moderate alteration of expression levels as effected by common genetic variation may cause a continuum of phenotypic changes. Alternatively, a more complex interplay between common and rare variation could be taking place such as higher tolerance to loss of function mutations in lower expressing haplotypes which could induce association with predicted expression.

We are finding that most trait associations are tissue specific; i.e. they are detected in a handful of tissues. However, we also find that expected tissues given known biology do not necessarily rank among the top enriched tissues. This suggests context specificity of the pathogenic mechanism; specific developmental stage or environmental conditions may be necessary to detect the regulatory event. On the other hand, we are detecting associations in unexpected tissues which suggests a sharing of regulation across multiple tissues / contexts or perhaps novel biology that takes place in these tissues. In either case, agnostic scanning of a broad set of tissues is necessary to discover these mechanisms.

### Software and Resources

We make our software publicly available on a GitHub repository: https://github.com/hakyimlab/MetaXcan. Prediction model weights and covariances for different tissues can be downloaded from PredictDB. A short working example can be found on the GitHub page; more extensive documentation can be found on the project’s [wiki](https://github.com/hakyimlab/MetaXcan/wiki) page. The results of MetaXcan applied to the 44 human tissues and a broad set of phenotypes can be queried on gene2pheno.org.

## Methods

### Summary-PrediXcan formula

Figure 1-B shows the main analytic expression used by Summary-PrediXcan for the Z-score (Wald statistic) of the association between predicted gene expression and a phenotype. The input variables are the weights used to predict the expression of a given gene, the variance and covariances of the markers included in the prediction, and the GWAS coefficient for each marker. The last factor in the formula can be computed exactly in principle, but we would need additional information that is unavailable in typical GWAS summary statistics output such as phenotype variance and sample size. Dropping this factor from the formula does not affect the accuracy of the results as demonstrated in the close to perfect concordance between PrediXcan and Summary-PrediXcan results on the diagonal of Figure 2-A.

The approximate formula we use is:

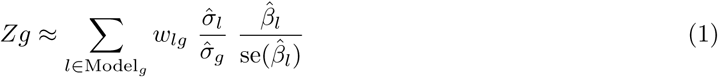

where

- *w*_*lg*_ is the weight of SNP *l* in the prediction of the expression of gene *g*,
- 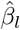 is the GWAS regression coefficients for SNP *l*,
- 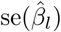 is standard error of 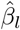,
- 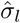 is the estimated variance of SNP *l*, and
- 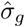 is the estimated variance of the predicted expression of gene *g*,
- dosage and alternate allele are assumed to be the same.

The inputs are based, in general, on data from three different sources:

- study set (e.g. GWAS study set),
- training set (e.g. GTEx, DGN),
- population reference set (e.g. the training set or 1000 Genomes).

The study set is the main dataset of interest from which the genotype and phenotypes of interest are gathered. The regression coefficients and standard errors are computed based on individual-level data from the study set or a SNP-level meta-analysis of multiple GWAS. Training sets are the reference transcriptome datasets used for the training of the prediction models (GTEx, DGN, Framingham, etc.) thus the weights *w*_*lg*_ are computed from this set. Training sets can also be used to generate variance and covariances of genetic markers, which will usually be different from the study sets. When individual level data are not available from the training set we use population reference sets such as 1000 Genomes data.

In the most common use scenario, users will need to provide only GWAS results using their study set. The remaining parameters are pre-computed https://github.com/hakyimlab/MetaXcan.

### Association enrichment

We display the enrichment for selected phenotypes in Supplementary Figure 5, measured as mean(*Z*^2^). For visualization purposes, we selected 25 phenotypes from different categories such as anthropometric traits, cardiometabolic traits, autoimmune diseases, and psychiatric conditions (please see figure caption for the list of selected phenotypes). The simple mean of *Z*^2^ for all gene-tissue pairs in a phenotype was taken.

### Derivation of Summary-PrediXcan Formula

The goal of Summary-PrediXcan is to infer the results of PrediXcan using only GWAS summary statistics. Individual level data are not needed for this algorithm. We will introduce some notations for the derivation of the analytic expressions of S-PrediXcan.

### Notation and Preliminaries

*Y* is the *n*-dimensional vector of phenotype for individuals *i* = 1, *n*. *X*_*l*_ is the allelic dosage for SNP *l*. *T*_*g*_ is the predicted expression (or estimated GREx, genetically regulated expression). *w*_*lg*_ are weights to predict expression 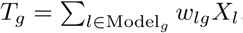, derived from an independent training set.

We model the phenotype as linear functions of *X*_*l*_ and *T*_*g*_

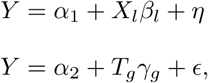

where *α*_1_ and *α*_2_ are intercepts, *η* and *∊* error terms independent of *X*_*l*_ and *T*_*g*_, respectively. Let 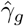 and 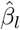 be the estimated regression coefficients of *Y* regressed on *T*_*g*_ and *X*_*l*_, respectively. 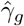 is the result (effect size for gene *g*) we get from PrediXcan whereas 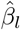 is the result from a GWAS for SNP *l*.

We will denote as 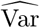 and 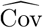 the operators that compute the sample variance and covariance, i.e.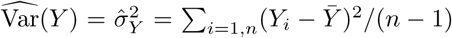 with 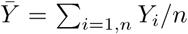. Let 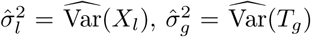 and 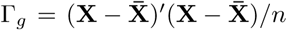, where **X**^′^ is the *p* × *n* matrix of SNP data and 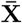 is a *n* × *p* matrix where column *l* has the column mean of **X**_*l*_ (*p* being the number of SNPs in the model for gene *g*, typically *p* ≪ *n*).

With this notation, our goal is to infer PrediXcan results (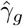 and its standard error) using only GWAS results (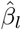 and their standard error), estimated variances of SNPs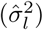, estimated covariances between SNPs in each gene model 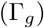, and prediction model weights *w*_*lg*_.

**Input**: 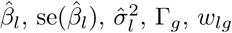. **Output**: 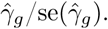

Next we list the properties and definitions used in the derivation:

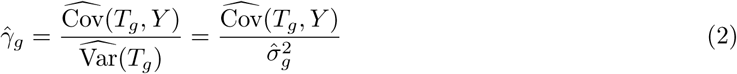

and

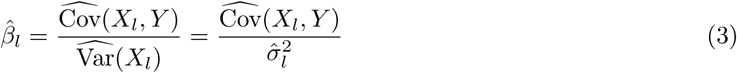

The proportion of variance explained by the covariate (*T*_*g*_ or *X*_*l*_) can be expressed as

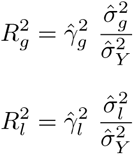

By definition

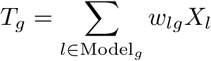

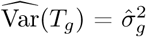 can be computed as

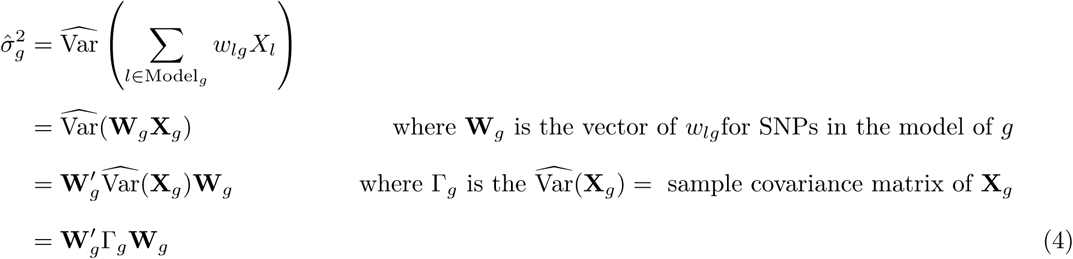

**Calculation of regression coefficient 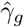**

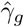 can be expressed as

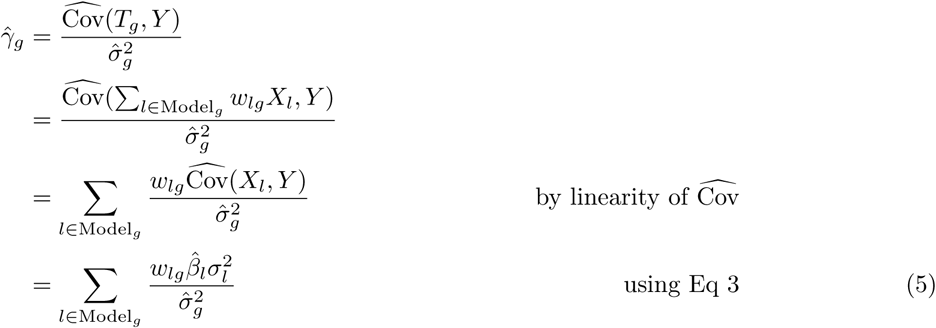

### Calculation of standard error of 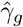

Also from the properties of linear regression we know that

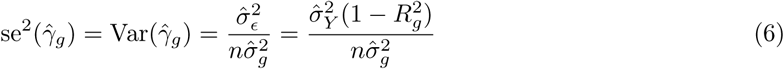

In this equation, 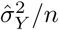 is not necessarily known but can be estimated using the analogous equation (6) for *β*_*l*_:

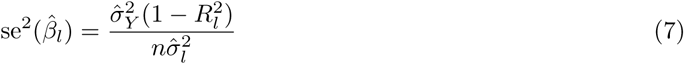

Thus:

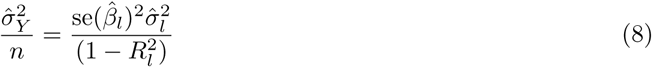

Notice that the right hand side of (8) is dependent on the SNP *l* while the left hand side is not. This equality will hold only approximately in our implementation since we will be using approximate values for 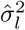, i.e. from reference population, not the actual study population.

### Calculation of Z-score

To assess the significance of the association, we need to compute the ratio of the estimated effect size 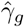 and standard error se(*γ*_*g*_), or Z-score,

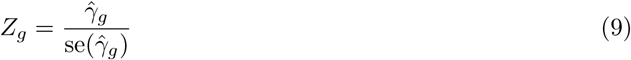

with which we can compute the p-value as 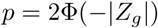 where Φ (.) is the normal CDF function.

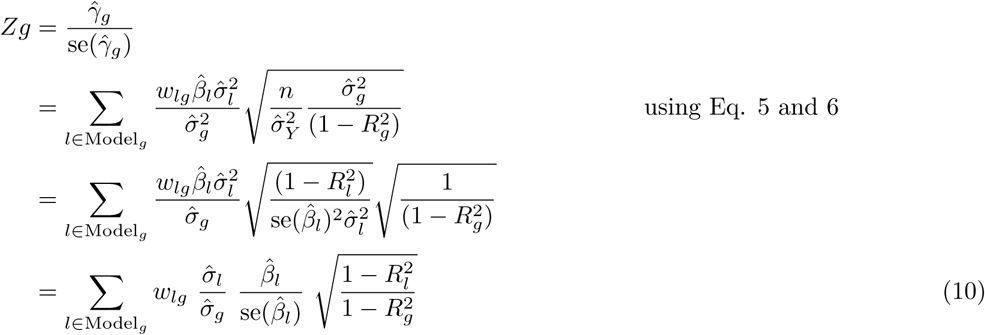

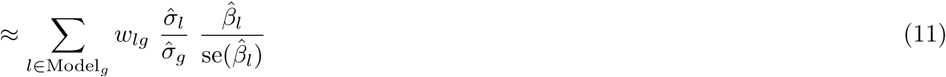

Based on results with actual and simulated data for realistic effect size ranges, we have found that the last approximation does not affect our ability to identify the association. The approximation becomes inaccurate only when the effect sizes are very large. But in these cases, the small decrease in statistical efficiency induced by the approximation is compensated by the large power to detect the larger effect sizes.

### Calculation of *σ*_*g*_ in reference set

The variance of predicted expression is computed using equation (4) which takes weights for each SNP in the prediction model and the correlation (LD) between the SNPs. The correlation is computed in a reference set such as 1000G or in the training set.

### Expression model training

To train our prediction models, we obtained genotype data and normalized gene expression data collected by the GTEx Project. We used 44 different tissues sampled by GTEx and thus generated 44 different tissue-wide models (dbGaP Accession phs000424.v6.p1). Sample sizes for different tissues range from 70 (Uterus) to 361 (Muscle-Skeletal). The models referenced in this paper make use of the GTEx Project’s V6p data, a patch to the version 6 data and makes use of improved gene-level annotation. We removed ambiguously stranded SNPs from genotype data, i.e. ref / alt pairs A / T, C / G, T / A, G / C. Genotype data was filtered to include only SNPs with MAF > 0.01. For each tissue, normalized gene expression data was adjusted for covariates such as gender, sequencing platform, the top 3 principal components from genotype data and top PEER Factors. The number of PEER Factors used was determined by sample size: 15 for n < 150, 30 for n between 150 and 250, and 35 for n > 250. Covariate data was provided by GTEx. For our analysis, we used protein-coding genes only.

For each gene-tissue pair for which we had adjusted expression data, we fit an Elastic-Net model based on the genotypes of the samples for the SNPs located within 1 Mb upstream of the gene’s transcription start site and 1 Mb downstream of the transcription end site. We used the R package glmnet with mixing parameter alpha equal to 0.5, and the penalty parameter lambda was chosen through 10-fold cross-validation.

Once we fit all models, we retained only the models q-value less than 0.05 [40]. For each tissue examined, we created a sqlite database to store the weights of the prediction models, as well as other statistics regarding model training. These databases have been made available for download at PredictDB.org.

### Online Catalog and SMR, COLOC, TWAS

Supplementary Table 4 shows the list of GWA / GWAMA studies we considered in this analysis. We applied S-PrediXcan to these studies using the transcriptome models trained on GTEx studies for patched version 6. For simplicity, S-PrediXcan only considers those SNPs that have a matching set of alleles in the prediction model, and adjusts the dosages (2 – dosage) if the alleles are swapped.

To make the results of this study broadly accessible, we built a Postgre SQL relational database to store S-PrediXcan results, and serve them via a web application http://gene2pheno.org.

We also applied SMR [16] to the same set of GWAMA studies, using the GTEx eQTL associations. We downloaded version 0.66 of the software from the SMR website, and ran it using the default parameters. We converted the GWAMA and GTEx eQTL studies to SMR input formats. In order to have SMR compute the colocalization test, for those few GWAMA studies where allele frequency was not reported, we filled in with frequencies from the 1000 Genomes Project [41] as an approximation. We also used the 1000 Genomes genotype data as reference panel for SMR.

Next we ran COLOC [18] over the same set of GWAMA and eQTL studies. We used the R package available from [CRAN](https://cran.r-project.org/web/packages/coloc/index.html). We used the Approximate Bayes Factor colocalization analysis, with effect sizes, their standard errors, allele frequencies and sample sizes as arguments. When the frequency information was missing from the GWAS, we filled in with data from the 1000 Genomes Project.

For comparison purposes, we have also included the results of the application of Summary-TWAS to 30 traits publicly shared by the authors [24].

### Comparison with TWAS

Formal similarity with TWAS can be made more explicit by rewriting S-PrediXcan formula in matrix form. With the following notation and definitions

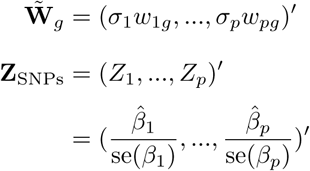

and correlation matrix of SNPs in the model for gene *g*

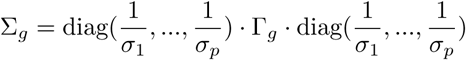

it is quite straightforward to write the numerator in (1) and (11) as

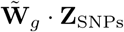

and in the denominator, the variance of the predicted expression level of gene *g*, as

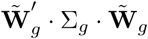

thus

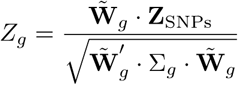

This equation has the same form as the TWAS expression if we use the scaled weight vector 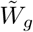 instead of *W*_*g*_. Summary-TWAS imputes the Z-score for the gene-level result assuming that under the null hypothesis, the Z-scores are normally distributed with the same correlation structure as the SNPs; whereas in S-PrediXcan we compute the results of PrediXcan using summary statistics. Thus, S-TWAS and S-PrediXcan yield equivalent mathematical expressions (after setting the factor 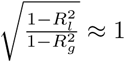).

### Summary-PrediXcan with only top eQTL as predictor

The S-PrediXcan formula when only the top eQTL is used to predict the expression level of a gene can be expressed as

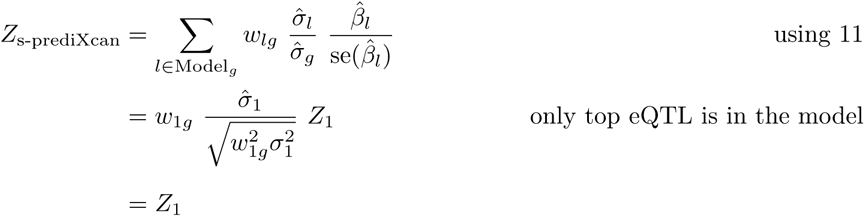

where *Z*_1_ is the GWAS Z-score of the top eQTL in the model for gene. Thus

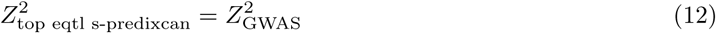

### Comparison with SMR

SMR quantifies the strength of the association between expression levels of a gene and complex traits with *T*_SMR_ using the following function of the eQTL and GWAS Z-score statistics:

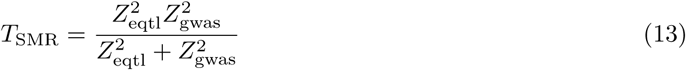

Here *Z*_eqtl_ is the Z-score (= effect size / standard error) of the association between SNP and gene expression, and *Z*_gwas_ is the Z-score of the association between SNP and trait.

This SMR statistic (*T*_SMR_) is not a 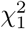 random variable as assumed in [16]. To prove this, we performed simulations following those described in [16]. We generated 10^5^ pairs of values for 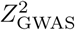 and 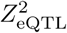. 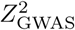 was sampled from a 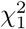 distribution. 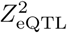 was sampled from a non-central 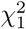 distribution with parameter *λ* = 29 (a value chosen to mimic results from [29], see [16]). Only pairs with eQTLs satisfying genome-wide significance (*p* < 5 × 10^−8^) were kept. We performed a QQ plot and observed deflation when comparing to random values sampled from a 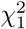 distribution (Supplementary Figure 5-B). This simulation was repeated 1000 times, and we observed a mean of *T*_SMR_ close to 0.93 (Supplementary Figure 5-C).

Only in two extreme cases, the chi-square approximation holds: when *Z*_eqtl_ ≫ *Z*_gwas_ or *Z*_eqtl_ ≪ *Z*_gwas_. In these extremes, we can apply Taylor expansions to find an interpretable form of the SMR statistic.

If *Z*_eqtl_ *≫ Z*_gwas_, i.e. if the eQTL association is much more significant than the GWAS association,

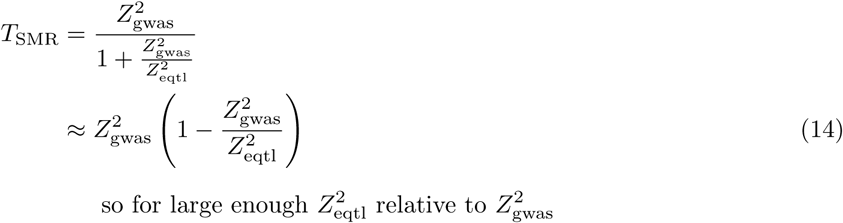

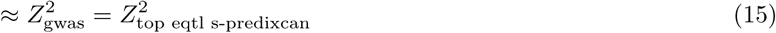

with the last equality from 12. Thus, in this case, the SMR statistic is slightly smaller than the (top eQTL based) S-PrediXcan *χ*_1_-square. This reduced significance is accounting for the uncertainty in the eQTL association. As the evidence for eQTL association grows, the denominator 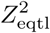 increases and the difference tends to 0.

On the other extreme when the GWAS association is much stronger than the eQTL’s, *Z*_eqtl_ ≪ *Z*_gwas_,

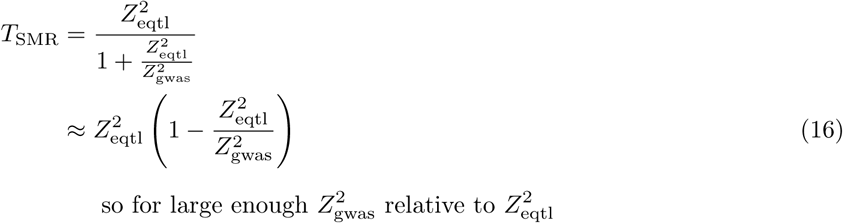

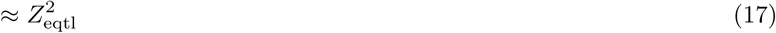

In both extremes, the SMR statistic significance is approximately equal to the less significant of the two statistics GWAS or eQTL, albeit strictly smaller.

In between the two extremes, the right distribution must be computed using numerical methods. When we look at the empirical distribution of the SMR statistic’s p-value against the GWAS and eQTL (top eQTL for the gene) p-values, we find the ceiling of the SMR statistic is maintained as shown in Figure 5-E and-F. Supplementary Figure 12 shows a comparison of colocalization proportions between SMR and PrediXcan.

### GERA imputation

Genotype files were obtained from dbGaP, and updated to release 35 of the probe annotations published by Affymetrix via PLINK [42]. Probes were filtered out that had a minor allele frequency of <0.01, were missing in >10% of subjects, or did not fit Hardy-Weinberg equilibrium. Subjects were dropped that had an unexpected level of heterozygosity (F >0.05). Finally the HRC-1000G-check-bim.pl script (from http://www.well.ox.ac.uk/∼wrayner/tools/) was used to perform some final filtering and split data by chromosome. Phasing (via eagle v2.3 [43]) and imputation against the HRC r1.1 2016 panel [44] (via minimac3) were carried out by the Michigan Imputation Server [45].

### GERA GWAS and MetaXcan Application

European samples had been split into ten groups during imputation to ease the computational burden on the Michigan server, so after obtaining the imputed.vcf files, we used the software PLINK [42] to convert the genotype files into the PLINK binary file format and merge the ten groups of samples together, while dropping any variants not found in all sample groups. For the association analysis, we performed a logistic regression using PLINK, and following QC practices from [14] we filtered out individuals with genotype missingness > 0.03 and filtered out variants with minor allele frequency < 0.01, missingness > 0.05, out of Hardy-Weinberg equilibrium significant at 1e-6, or had imputation quality < 0.8. We used gender and the first ten genetic principal components as obtained from dbGaP as covariates. Following all filtering, our analysis included 61,444 European samples with 7,120,064 variants. MetaXcan was then applied to these GWAS results, using the 45 prediction models (GTEx and DGN).

## Acknowledgments

### Grants

We acknowledge the following US National Institutes of Health grants: R01MH107666 (H.K.I.), T32 MH020065 (K.P.S.), R01 MH101820 (GTEx), P30 DK20595 (Diabetes Research and Training Center), F31 DK101202 (J.M.T.), P50 DA037844 (Rat Genomics), P50 MH094267 (Conte). H.E.W. was supported in part by start-up funds from Loyola University Chicago.

The Genotype-Tissue Expression (GTEx) Project was supported by the Common Fund of the Office of the Director of the National Institutes of Health. Additional funds were provided by the NCI, NHGRI, NHLBI, NIDA, NIMH, and NINDS. Donors were enrolled at Biospecimen Source Sites funded by NCI SAIC-Frederick, Inc. (SAIC-F) subcontracts to the National Disease Research Interchange (10XS170), Roswell Park Cancer Institute (10XS171), and Science Care, Inc. (X10S172). The Laboratory, Data Analysis, and Coordinating Center (LDACC) was funded through a contract (HHSN268201000029C) to The Broad Institute, Inc. Biorepository operations were funded through an SAIC-F subcontract to Van Andel Institute (10ST1035). Additional data repository and project management were provided by SAIC-F (HHSN261200800001E). The Brain Bank was supported by a supplements to University of Miami grants DA006227 & DA033684 and to contract N01MH000028. Statistical Methods development grants were made to the University of Geneva (MH090941 & MH101814), the University of Chicago (MH090951, MH090937, MH101820, MH101825), the University of North Carolina-Chapel Hill (MH090936 & MH101819), Harvard University (MH090948), Stanford University (MH101782), Washington University St Louis (MH101810), and the University of Pennsylvania (MH101822). The data used for the analyses described in this manuscript were obtained from dbGaP accession number phs000424.v6.p1 on 06/17/2016.

This work was completed in part with resources provided by the University of Chicago Research Computing Center, Bionimbus [46], and the Center for Research Informatics. The Center for Research Informatics is funded by the Biological Sciences Division at the University of Chicago with additional funding provided by the Institute for Translational Medicine, CTSA grant number UL1 TR000430 from the National Institutes of Health.

